# Examination of the role of mutualism in immune evasion

**DOI:** 10.1101/2024.03.02.583098

**Authors:** Lucie Gourmet, Simon Walker-Samuel, Parag Mallick

## Abstract

Though the earliest stages of oncogenesis, post initiation, are not well understood, it is generally appreciated that a successful transition from a collection of dysregulated cells to an aggressive tumor requires complex ecological interactions between cancer cells and their environment. One key component of tumorigenesis is immune evasion. To investigate the interplay amongst the ecological behaviour of mutualism and immune evasion, we used a computational simulation framework. Sensitivity analyses of the growth of a virtual tumor implemented as a 2D-hexagonal lattice model suggests tumor survival depends on the interplay between growth rates, mutualism and immune evasion. In 60% of simulations, cancer clones with low growth rates, but exhibiting mutualism were able to evade the immune system and continue progressing suggesting that tumors with equivalent growth rates and no mutualism are more likely to be eliminated than tumors with mutualism. Tumors with faster growth rates showed a lower dependence upon mutualism for progression. Geostatistical analysis showed decreased spatial heterogeneity over time for polyclonal tumours with a high division rate. Overall, these results suggest that in slow growing tumors, mutualism is critical for early tumorigenesis.

## Introduction

The earliest genetic aberrations in oncogenes and tumor supressors leading to transformation of a cell from healthy to malignant have been extensively studied. Additionally, it is generally established that critical cancer hallmarks such as genome instability, immune evasion and angiogenesis^1^ are required for tumor formation. However, the earliest stages of a tumor, including the transition from partially transformed collections of cells^2^ to a malignant tumor are not well understood. The traditional clonal evolution hypothesis implies that cancer originates from one cell^3^ which grows into a fully formed tumor. However, multiple studies have shown evidence of early cancer polyclonality^4^ suggesting that tumor formation may be a more complex process involving intercellular interactions and interplay between early cancer cells and the emerging tumor microenvironment. Because of the multiplicity of cell clones types, architectures and behaviours involved in tumorigenesis, cancer can be viewed as an ecosystem.

Recent studies^5^ have highlighted the role of ecological behaviours in later stage cancers and also emphasized the role that tumor ecology plays in both progression and therapeutic response^6^. These studies approach cancer growth from a population dynamics perspective in which different cell types compete for resources and survival and demonstrated the existence of cooperative behaviours such as mutualism and commensalism amongst cancer cells. Cooperation can be defined as mutualism, where both cancer cells benefit from interacting, or commensalism where only one of them benefits. One might also describe immune-cancer interactions as predation as interactions between immune cells and cancer cells typically lead to death of targeted cancer cells. Mechanistically, ecological behaviors amongst cancer cells are driven by either molecular or physical interactions amongst cells. Two main types of chemical interactions are relevant for cancer cooperation: juxtacrine and paracrine signalling. Juxtacrine signalling occurs at short distances since it is dependent on cell contact to pass signalling molecules directly between cells^7^. In contrast, paracrine signalling is the diffusion of signalling molecules from one cell to another but does not require cell-cell contact^7^. Paracrine signalling was shown to increase proliferation: for example, IL-6 promotes proliferation between heterogeneous breast cancer subclones^8^.Conversely, paracrine signalling of TGF-β was shown to promote immune suppression^9^. Though there has been substantial study of intercellular communication and the molecular and cellular consequences thereof, the potential ecological impacts of that communication have not been as well understood.

Multiple studies suggest that cooperation is essential for cancer evolution. Early cancer formation in mice were found to be polyclonal when skin papillomas were induced by chemicals^10^. This finding was the first to suggest that interaction between cells is necessary for carcinogenesis rather than the proliferation of a single clone. Moreover, the interleukin-6 family cytokines were shown to be involved in tumour heterogeneity and required for sustained tumour growth^11^. This study found that minor subclones (which make up to 10% of the tumour population) were not outcompeted because they secrete Il-11 and FIGF. This is an advantage because they induce metastasis by modulating the immune system and blood vessels. Thus, the presence of such clones enables angiogenesis, one of the hallmarks of cancer, which benefits the whole tumour. Interclonal cooperation was also shown to be critical for tumour maintenance in a mouse models of breast cancer^12^. The two clones (Wnt^high^HRAS^wt^ and Wnt^low^HRAS^mut^) present in the mammary tumour are essential for cancer as they form tumours only when combined. The interplay between cancer cells at an early stage is therefore important for cancer development.

Even if early tumours arise, they can also be removed by the immune system or by cell competition. Cancer immunoediting occurs in 3 stages: elimination, equilibrium and escape^13^. Elimination represents the process of immunosurveillance while equilibrium is the selection of immune resistant clones. Finally, immune escape means that the cancer is no longer controlled by the immune system and can spread. Immune escape is an important aspect of cancer because it impacts prognosis. For example, patients with pancreatic cancer showed higher survival when subjected to strong immune editing^14^. Nevertheless, clonal cooperation can hinder immune predation as clonal cooperation was shown to activate pro-proliferative signalling pathways such as JNK and IL-6^15^. Cell contact–mediated mechanisms can also promote tumour growth as breast cancer cell lines with PIK3CA mutant cells were able to induce the proliferation of quiescent HER2 mutant through fibronectin interactions^16^.

We hypothesize that early cell-cell interactions determine the effectiveness of carcinogenesis with immune predation and clonal cooperation acting as opposing forces. More specifically we test the hypothesis that mutalism can be a mediator of successful immune escape and examined the three-way interactions between cell’s basal growth rate, mutualism and immune cell predation. To test these hypotheses we extended the model of Boyce and Mallick^17^ (**Fig 1**). We define the difference in growth trajectory between tumors within and without immune predation as K and estimate K using the Kolmogorov-Smirnov statistic.

**Figure 1:**
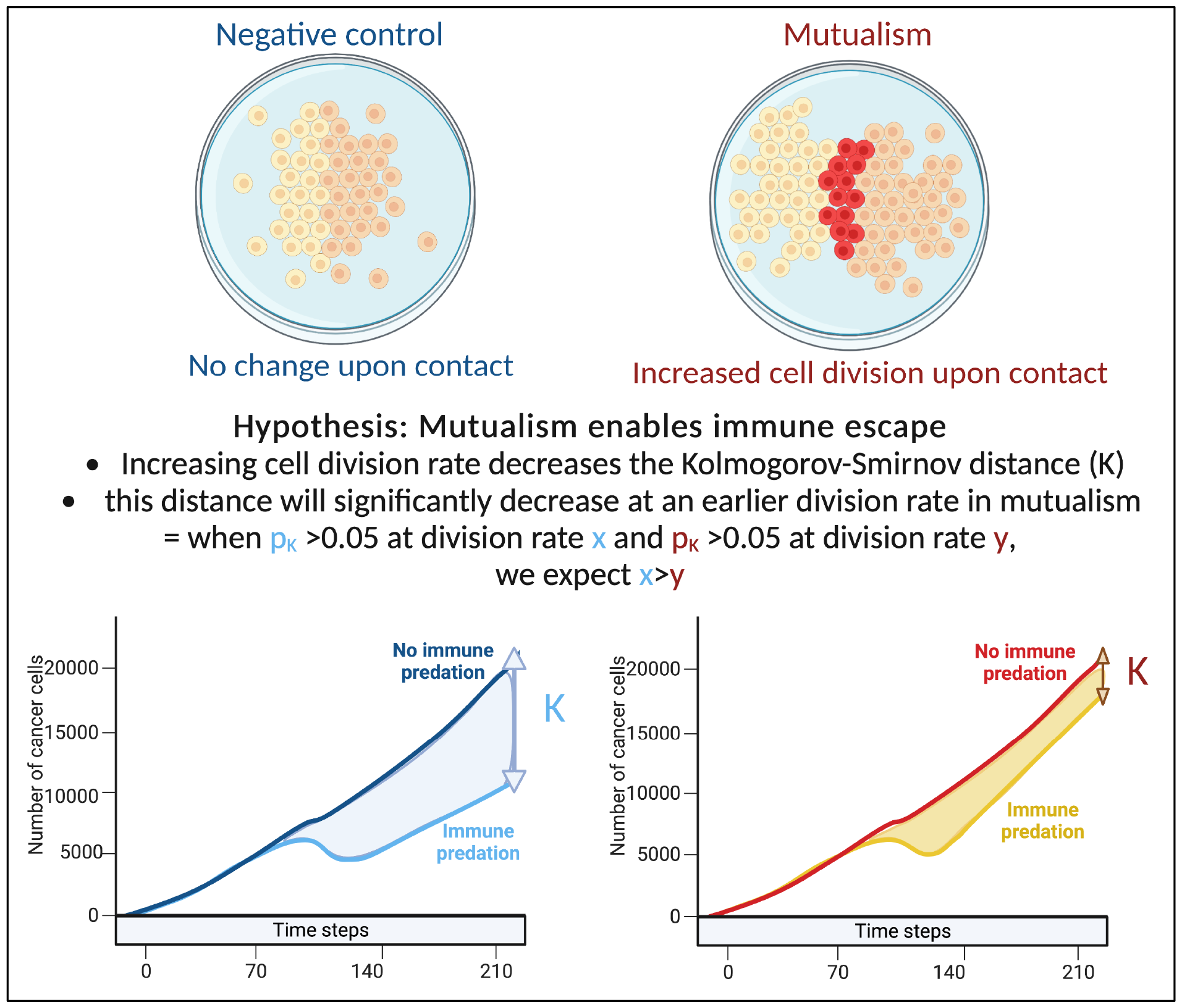
Explanation of our hypothesis and methodology.

## Methods

### Agent-based modelling

Here we briefly describe the modelling framework used for our simulation studies. We note that the additional detail about the framework and the geostatistical measures used herein are explained in more detail in our previous paper^17^. Our tumour growth and ecology model is implemented as a lattice-based cellular automaton which considers two-dimensional space as a hexagonal grid where each position has six direct neighbours with which it can interact. The model is initialized with an initial number of cancer cells, intrinsic oncoprotein-expression related birth and death rates for each cell, rules for ecological interactions and the dimensions of the canvas. Upon cell division, the cellular automaton looks for space to divide and chooses the position closest to the model centre to add a new cell. It keeps track of the number and types of agents along with the dimensions of the canvas. Because cells that are spatially proximal tend to have correlated expression, oncoprotein initialization is dependent on the expression of the neighbours of a given cell. To achieve spatial autocorrelation, we utilize a Markov Chain Monte Carlo sampling strategy that incorporates the mean oncoprotein expression of a cellar neighbourhood^17^.

### Ecological behaviours

To model mutualism, we initialize two separate clones on the canvas, each with cancer cells and cancer stem cells, and the tumours are allowed to grow^17^. When we model mutualism, if a cancer cell from the first clone contacts a cell from the second clone, a mutualistic interaction occurs and both cells have an increase in oncoprotein expression and gain a fitness benefit and an increase in instantaneous proliferation rate of 1.25. This increase is not compound and is only allowed to happen once. In a non-mutualistic setting, there is no proliferation advantage upon contact between cells of different clonal origin.

We model immune interactions in the tumour microenvironment via a predation model. Here, we initialized the tumour model with cancer cells and cancer stem cells in the centre of the model canvas and initialize a perimeter of immune cells encircling the tumour^17^. Immune cells are inactive until the tumour reaches a cancer cell count threshold of 3000, upon which the immune cells are activated. While the tumour continues to grow, immune cells begin to move with preference toward the weighted centre of the cancer cells. When an immune cell neighbours a cancer cell it will target and attach to the cancer with its initialized target probability of 0.99. If an immune cell is already targeting a cancer cell, it will kill that cell with its initialized killing probability of 0.99. Immune cells can only target one cancer cell at a time and targeting and killing occur at different steps of the model. In our simulations, we combined the predation model with the mutualism model such that our composite model includes the immune system encircling two cancer clones.

### Geostatistics

We calculate three global statistics at each step that describe how the spatial heterogeneity of oncoprotein in the virtual tumor^17^. To quantify general clustering of oncoprotein at the global level we use the Getis-Ord general G statistic. The G statistic is bounded between 0, indicating clustering of low values, and 1 indicating clustering of high values. To quantify spatial heterogeneity, we utilize two statistics: Moran’s I statistic and Geary’s C. The value of I is between −1 for perfectly dispersed (low autocorrelation) data and +1 for perfectly clustered (high autocorrelation) data. In contrast, Geary’s C is inversely related to Moran’s I and can take on values between 0 with perfect clustering (high autocorrelation) and an undefined upper bound for increased dispersion (low autocorrelation).

### Comparing cancer growth across different division rates

We examined the impact of cellular growth rate on tumor growth trajectory. As there are non-linear and spatial interactions in the model, tumor growth rate is not simply a function of cellular growth rate. For mutualism to occur, the two clones need to come into contact. Our sensitivity analysis examined division rates between 0.01 to 0.1 cell divisions per hour. This range of rates includes the experimentally estimated growth rates of non-small-cell lung cancer cells in 2D cell cultures that have been estimated to have a division rate close to 0.0461 divisions per hour^18,19^. To compare the tumor growth trajectories we used the Kolmogorov-Smirnov distance: when the associated p value is above 0.05, we consider that the number of cancer cells with/without predation are statistically similar. If the number of cancer cells with/without immune predation is similar at a lower division rate in the mutualistic setting compared to no mutualism, we can conclude that mutualism provides a significant advantage (**Fig 1**).

### Randomisation analysis

We estimated the number of mutualistic cells with advantageous proliferation (mutual cells with oncoprotein expression superior to 4 as shown in red in SuppFig3) and the number of cancer cells. We then used this ratio as a rate of random mutualism: for example, we found that on average 3% of all cancer cells are mutualistic cells dividing with a proliferation advantage. Thus in this general case, instead of mutualism upon clonal contact, we set the condition np.random.uniform() > 0.97 for all cells, which means that there is a 97% likelihood the cell will be a regular cancer, and a 3% chance it will become randomly mutualistic. To ensure that the random analysis reflects the mutualism model, we update the rate of random mutualism at every step accordingly to the rate of the mutualism model (**SuppFig 2**).

## Results

### Differential growth trajectory of mutualistic and non-mutualistic tumors

To visualise the impact of immune predation and mutualism on tumor growth, we performed a sensitivity analysis across a range of cellular growth rates and plotted tumor growth trajectory curves. At cell division rates below .03 divisions per hour, cells from the two clonal populations do not interact given their initial distance and are unable to benefit from mutualism. For each cellular growth rate, we simulated 4 different scenarios: mutualism with/without immune predation (in orange and red respectively) and no mutualism with/without immune predation (in turquoise and blue respectively). We observed that the mutualistic scenarios did not yield the highest number of cancer cells (**Fig 2A**). However, this heavily depended on the cancer cell division rate. Looking at a division rate of 0.08 division per hour, mutualism eventually displayed the highest number of cancer cells (**Fig 2B**). Interestingly, at even higher growth rates (for example 0.09), the mutualistic and no mutualistic curves seem to completely overlap (**SuppFig 1**). We therefore observe that when the cellular division rate is high enough, mutualism is not as impactful on overcoming the impact of immune-mediated cell death.

**Figure 2:**
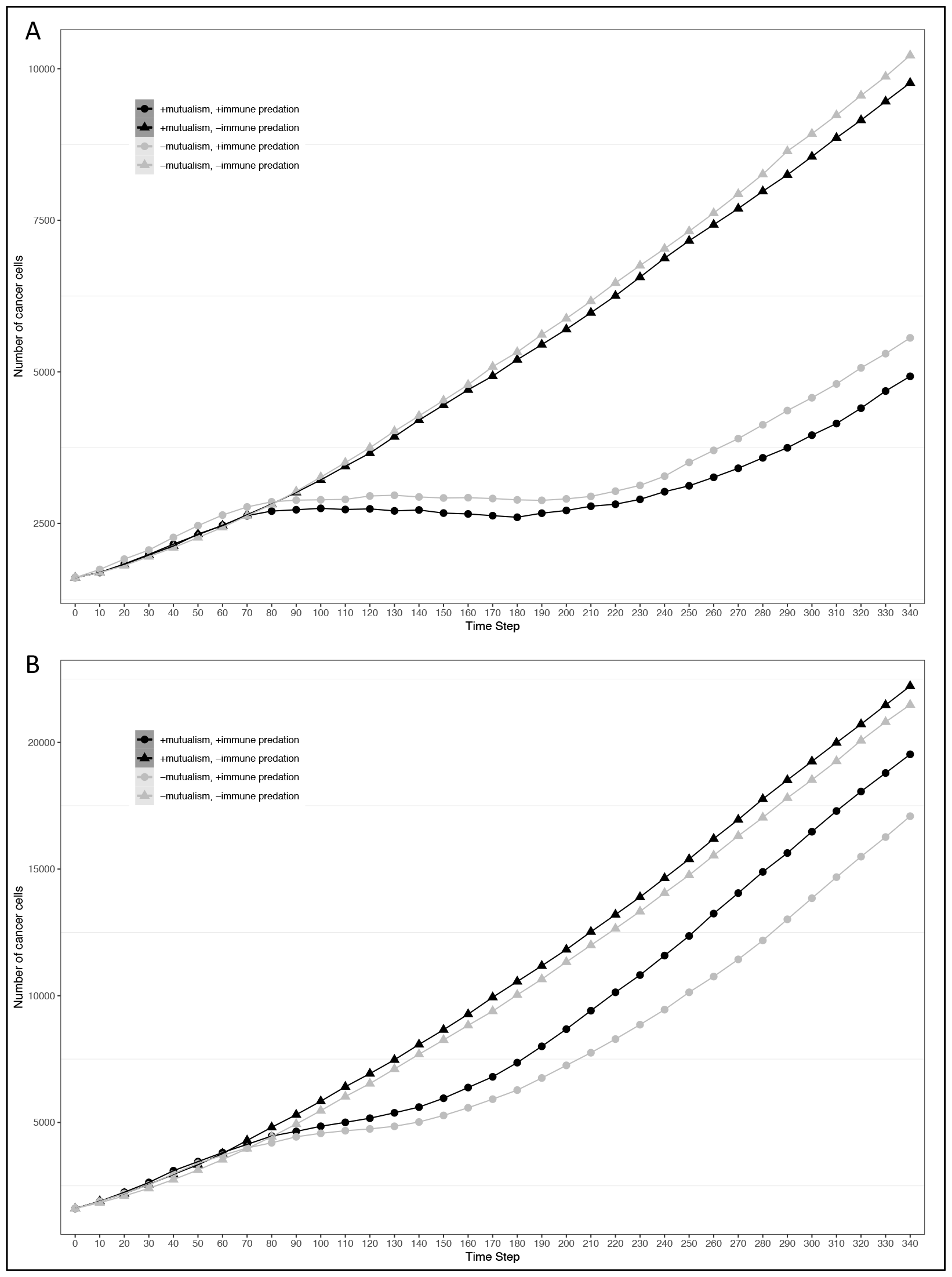
Growth curves representing the number of cancer cells depending on different conditions (the presence immune predation and/or mutualism). The x axis represents time steps while the y axis is the number of cancer cells. A: 0.04 division per hour. B: 0.08 division per hour.

### Mutualism enables compensation for immune predation

To assess the impact of mutualism on immune predation, we did 10 simulations for each cellular division rate. Across the different cellular division rates, 6 out of 10 of the mutualistic virtual tumors had higher growth trajectories than the non-mutualistic virtual tumors. One simulation showed no difference between mutualism and no mutualism. In 3 non-mutalistic virtual tumors had higher growth trajectories than the mutualistic virtual tumors. The results from all the simulations are summarised in a violin plot representing the Kolmogorov-Smirnov distance versus the division rate (**Fig 3**). Mutualism starts when the division rate is high enough (around 0.03) to enable contact between the clones before one of them is removed by the immune system. Interestingly, already at this division rate (0.03), we can observe that mutualism is advantageous.

**Fig 3:**
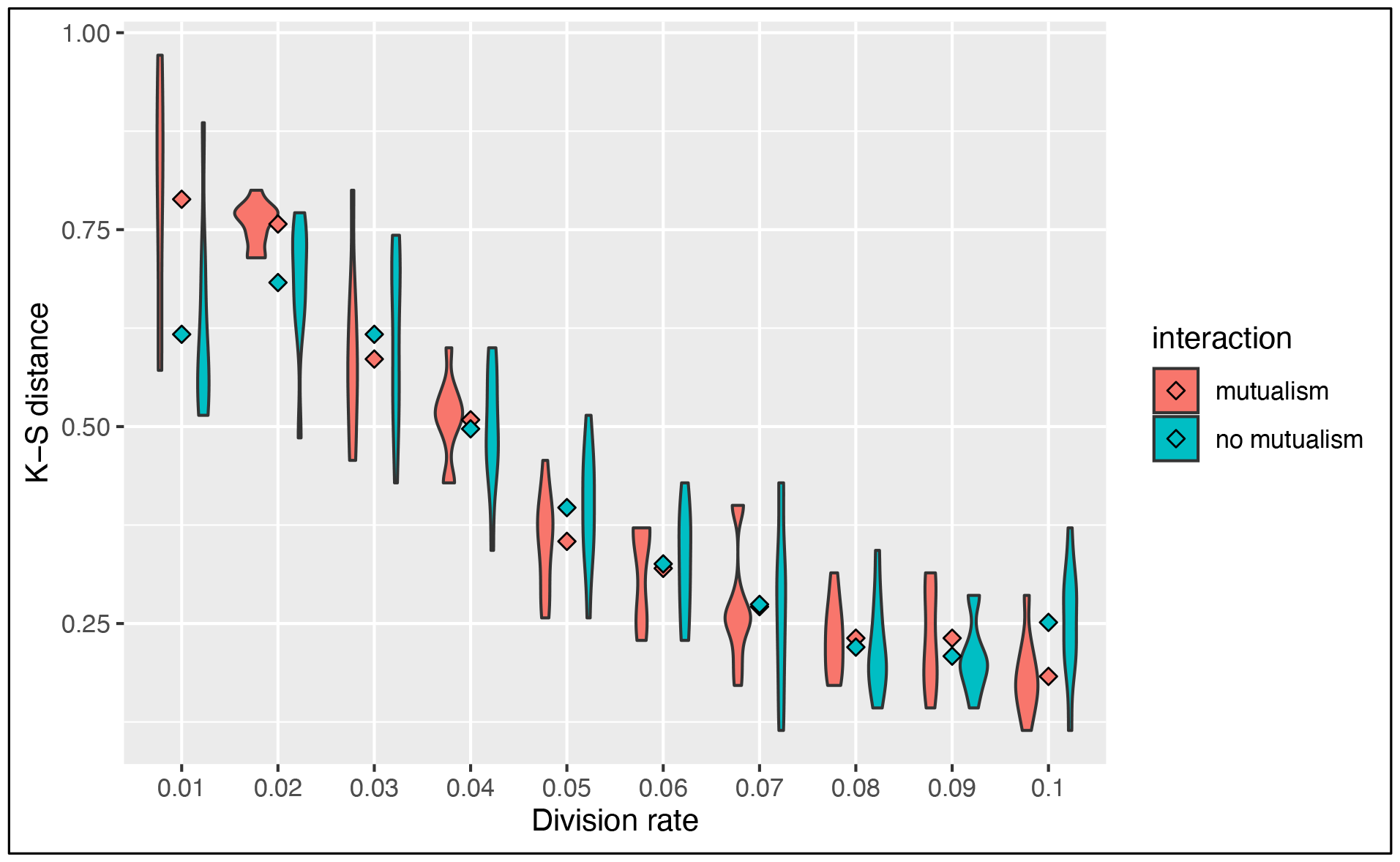
Summary of sensitivity analysis of tumor growth trajectory in the presence of immune predation. The Kolmorgorov-Smirnov distance is used to compare mutualism (in red) and no mutualism (in blue) across different division rates. The lower the Kolmogorov-Smirnov distance, the more similar the number of cancer cell is between immune predation and no immune predation.

### Mutualism rather than just the number of clones promotes cancer survival

In the previous paper, Boyce and Mallick focused on immune predation in the case of one cancer clone. Here, we analyse the geostatistics upon the presence of two distinct clonal populations (**Fig 4**). We notice that N, the delta number of cancer cells, becomes constant over time indicating that the tumour neither grows nor shrinks. Compared to the monoclonal predation model, the geostatistics vary considerably. Geary’s C is continually increasing, reflecting the fact that spatial heterogeneity increases over time. In contrast, the other two metrics stay constant unlike the monoclonal predation model in which there is an eventual increase. Having two clones instead of one promotes heterogeneity, as we may expect, and seems to lead to more stable geostatistics overall.

**Fig 4:**
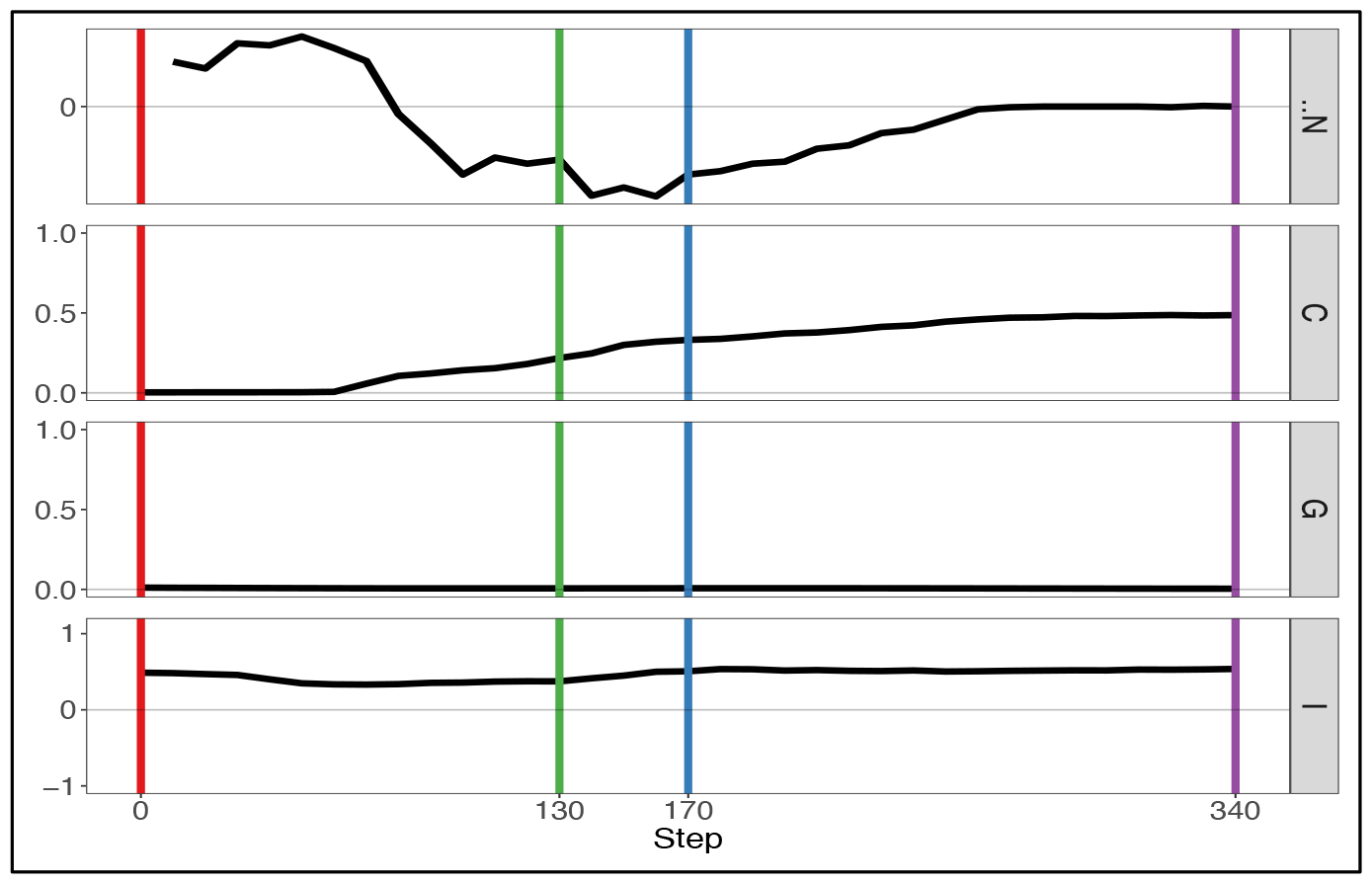
Geostatistics of two cancer clones without mutualism and with immune predation. Metrics from top to bottom: N the delta number of cancer cells from the previous step, Geary’s C, Getis-Ord G, Moran’s I.

If we enable mutualistic interactions in the predation model, we see little change from our previous results. The main difference is the fact that N, the delta number of cancer cells, continues to increase instead of staying constant at a certain point (**Fig 5**). If we compare our results of our simulation with the simulation representing mutualism only, we observe more similarities. The metrics Getis-Ord G and Moran’s I follow the same pattern, thereby indicating that mutualism determines spatial heterogeneity more than immune predation. Therefore, this result already supports observation that mutualism outcompetes the immune system, as the output of the simulation is closer to the mutualistic results.

**Fig 5:**
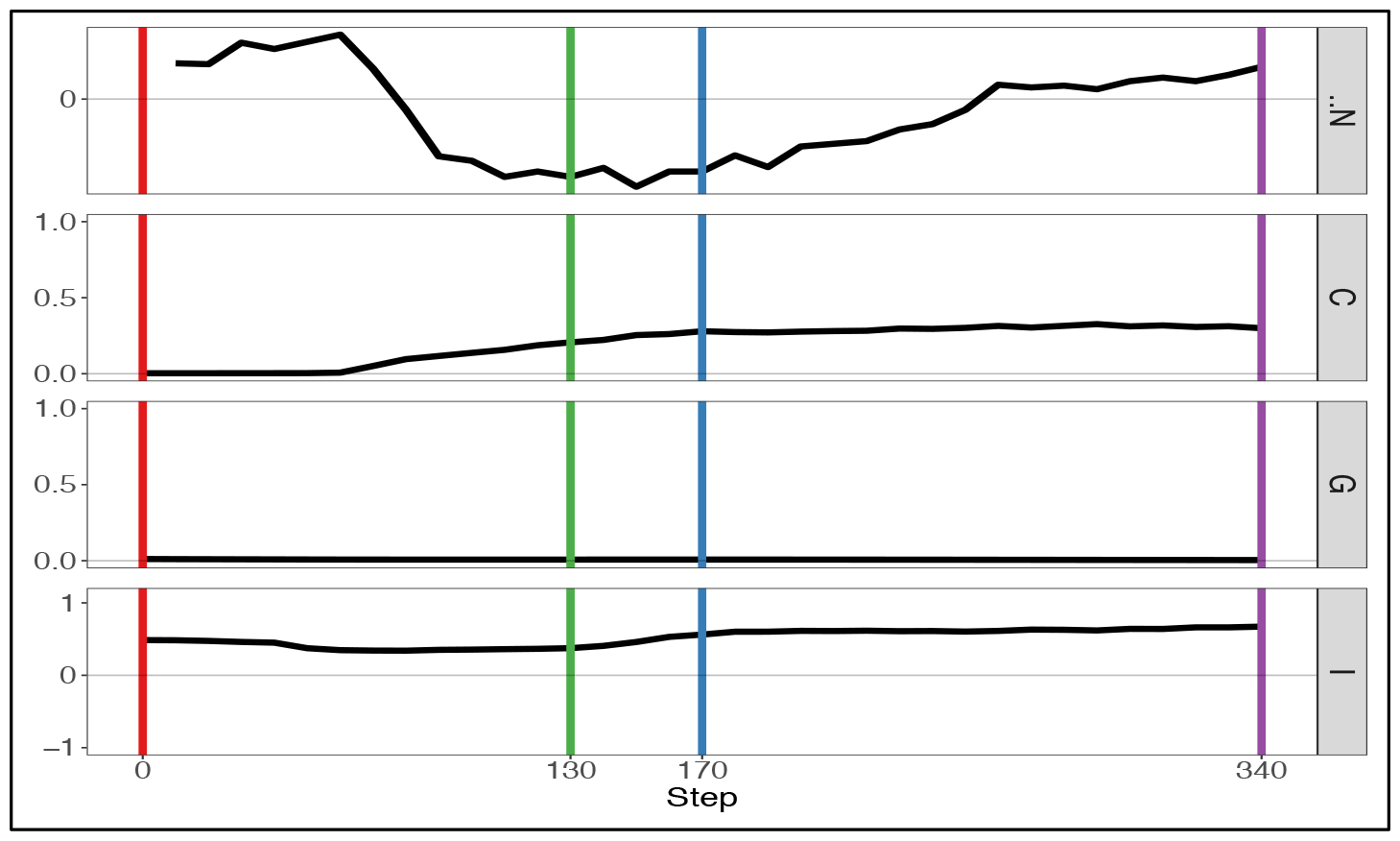
Geostatistics of two cancer clones with mutualism and immune predation. Metrics from top to bottom: N the delta number of cancer cells from the previous step, Geary’s C, Getis-Ord G,Moran’s I.

### Polyclonal tumours with high division rate also become homogeneous over time

With this simulation, we explore what happens to cancer clones which have a high division rate (which can be considered malignant) but also what happens in the long term. Therefore, we used a higher division rate (0.1) and a longer timeframe (490 steps instead of 340 as showed in **Fig 6**). We notice that there is much more variation in the geostatistical values due to the increase division rate. For instance, Geary’s C increases and decreases significantly, leading to a homogeneous tumour. It seems that increasing the potential for mutualism increases spatial heterogeneity, but the level of heterogeneity remains low eventually. Moreover, the delta number of cancer cells N starts to decrease as the tumour is filling all the available space left.

**Fig 6:**
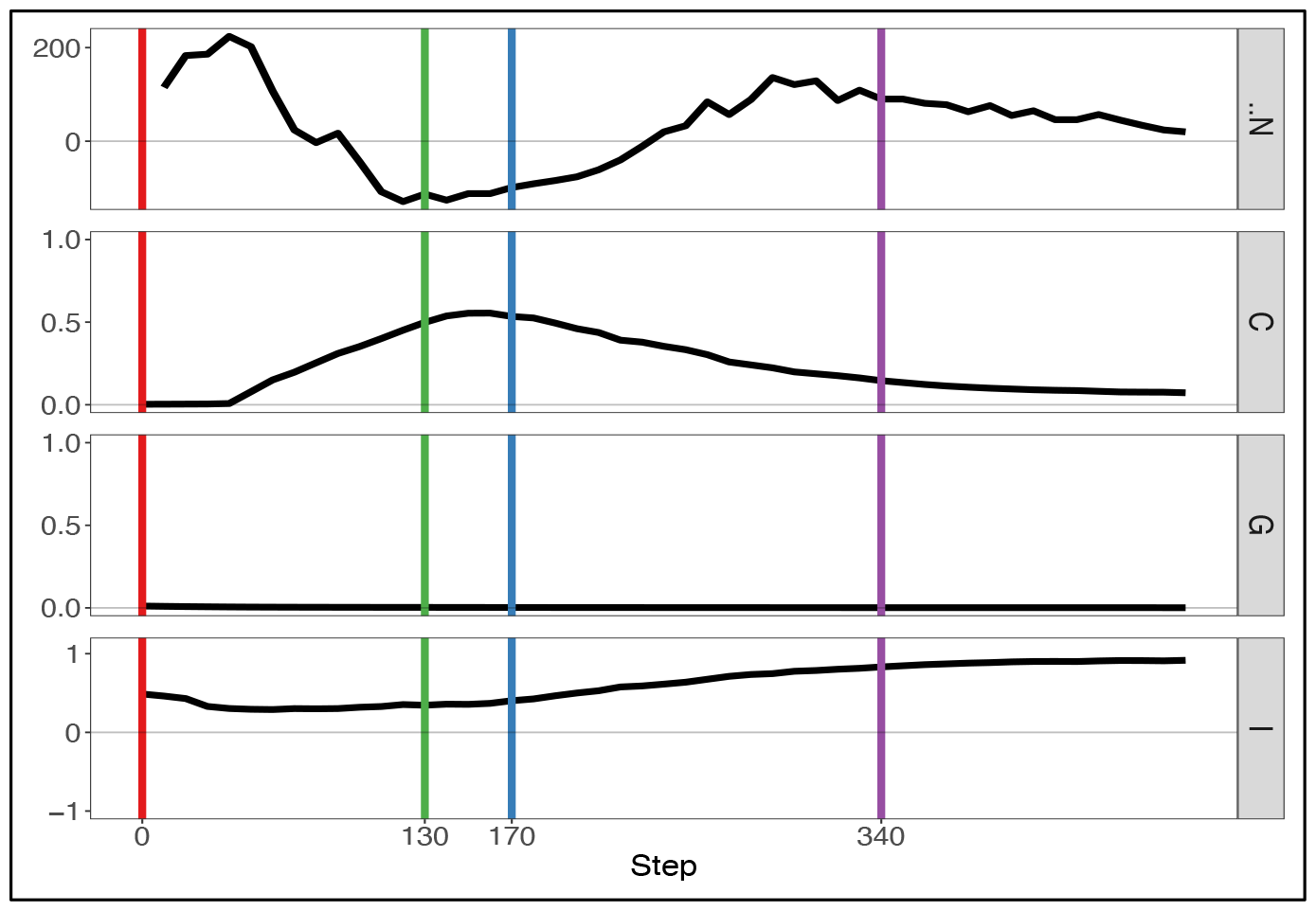
Geostatistics of two malignant (division rate 0.1) clones with mutualism and immune predation. Metrics from top to bottom: N the delta number of cancer cells from the previous step, Geary’s C, Getis-Ord G,Moran’s I.

## Discussion

By analysing growth curves, we observed that mutualism provides a tumor growth advantage at high division rates (**Fig 2**). We observe that in the majority of simulations mutualism makes a statistical difference to enable tumors to compensate for immune predation (**Fig 3**). When there is immune predation, having two clones instead of one has important consequences regarding cancer evolution. Indeed, we show in Figure 4 that it promotes spatial heterogeneity. Adding mutual interactions to this simulation increases N, the delta number of cancer cells, implying that mutualism enables sustained tumour growth. Without mutualism, the tumour seems to be contained (even though it is not completely removed) but mutualism seems necessary for continuing further cancer development (**Fig 5**). Increasing the division rate does increase drastically spatial heterogeneity, but the tumour becomes homogeneous eventually (**Fig 6**). We note that this is not due to having a larger number of cells at tumor initiation. Additionally, this behavior would not be observed with a fixed number of cells randomly having higher growth rates. In fact, we discovered that when there is immune predation, mutualism is more advantageous than random mutualism (**SuppFig 2**).

We analysed the cases in which no mutualism was more advantageous than mutualism and we found that cell distribution seems to impact whether immune predation is outcompeted. The location and pattern of mutualistic cells varied between the cases that support our hypothesis and those which do not (**SuppFig 3**). From a qualitative point of view, mutualism did not seem to help outcompeting immune predation when it was restricted to a specific part of the clone (**SuppFig 3**). Other factors to consider are the way in which the tumours merge and the distribution of immune cells around the tumour. The overall spatial composition of the tumour and its environment seems to be key for its evolution.

There are a few limitations to approach used in this study, including the fact that the model employed is a 2D on-lattice model which does not incorporate 3-dimensional or continuum aspects of the tumour (the diffusion of molecules, angiogenesis, cell morphology, long range communication…). As our model is dominantly a cell scale model, it does not have a specific set of molecular pathways responsible for the change in cellular behaviours. If it did, we could have modelled signalling pathways responsible of clonal cooperation instead of looking at contact-mediated mutualism. If we had used an off-lattice model, which is based on physical interactions, our results would be more realistic. Besides, the outcome of our model is mainly defined by the balance between the rate of cell division and cell death, but we investigated growth kinetics across multiple proliferation rates to overcome this shortfall. Moreover, the parameters of our model come from experiments involving non-small lung cancer cell line meaning that our results are relevant to this specific cell type. In the end, it is difficult to generalise the results of our simulation to real tumours because of the limitations mentioned above. Nevertheless, using our agent-based model was advantageous because we modelled an event difficult to measure and it was not computationally expensive.

In the future, by understanding the mechanisms underlying mutualism (mutations, down/upregulation of genes…), it may be possible to stop the development of cancer not through cytotoxic or cytostatic therapies, but instead by therapies that target mutualism specifically. Finding signatures of cancer cooperation could help us make sense of early carcinogenesis. Moreover, it could have a clinical impact if we not only investigate experimental models but also unravel interactions in patient samples. The key conclusion from our study is that mutualism will increase cancer proliferation in a manner that enables slower growing cells to evade the immune system until they reach a critical threshold in which the effective tumor growth rate exceeds the cell-killing capacity of tumor-infiltrating immune system. Characterising cooperation in cancer at an early stage is therefore essential to prevent cancer formation. A therapy strategy would be to selectively remove mutualistic cells, known as “common gooder” cells which have paracrine influence, to make the tumour collapse^20^.

Furthermore, our hypothesis makes a few assumptions such as the fact that cancer cooperation necessarily leads to an increase of proliferation. As we already mentioned, there are various examples of mutualism where collaboration involves sharing cancer hallmarks rather than a direct consequence on cell division. Interestingly, cancer cells have recently been shown to survive immune predation by hiding inside other cancer cells^21^. This cell-in-cell formation implies that only the internal hidden cancer cell survives but this mechanism provides resistance against T cells attacks and chemotherapies. We also consider that mutualistic cells have a high frequency, even though they may be outcompeted by a “free rider” clone. Nevertheless, because we focus on early carcinogenesis (a time during which tumours are more easily removed by immune predation), the later stages of cancer development do not matter that much. We also consider a tumour size threshold after which immune predation cannot outcompete the tumour anymore, which may not be the case. However, a study supports our hypothesis as its model predicts the existence of an antigen diversity threshold level beyond which T cells fail at controlling heterogeneous tumours^22^. A recent study showed that a specific clone can suppress anti-cancer CD8 T cell responses and protect other clones from the immune system. It would be interesting to further investigate the gene signature responsible for this immunotherapy resistance and poor survival in breast cancer patients.

---

**SuppFig1:**
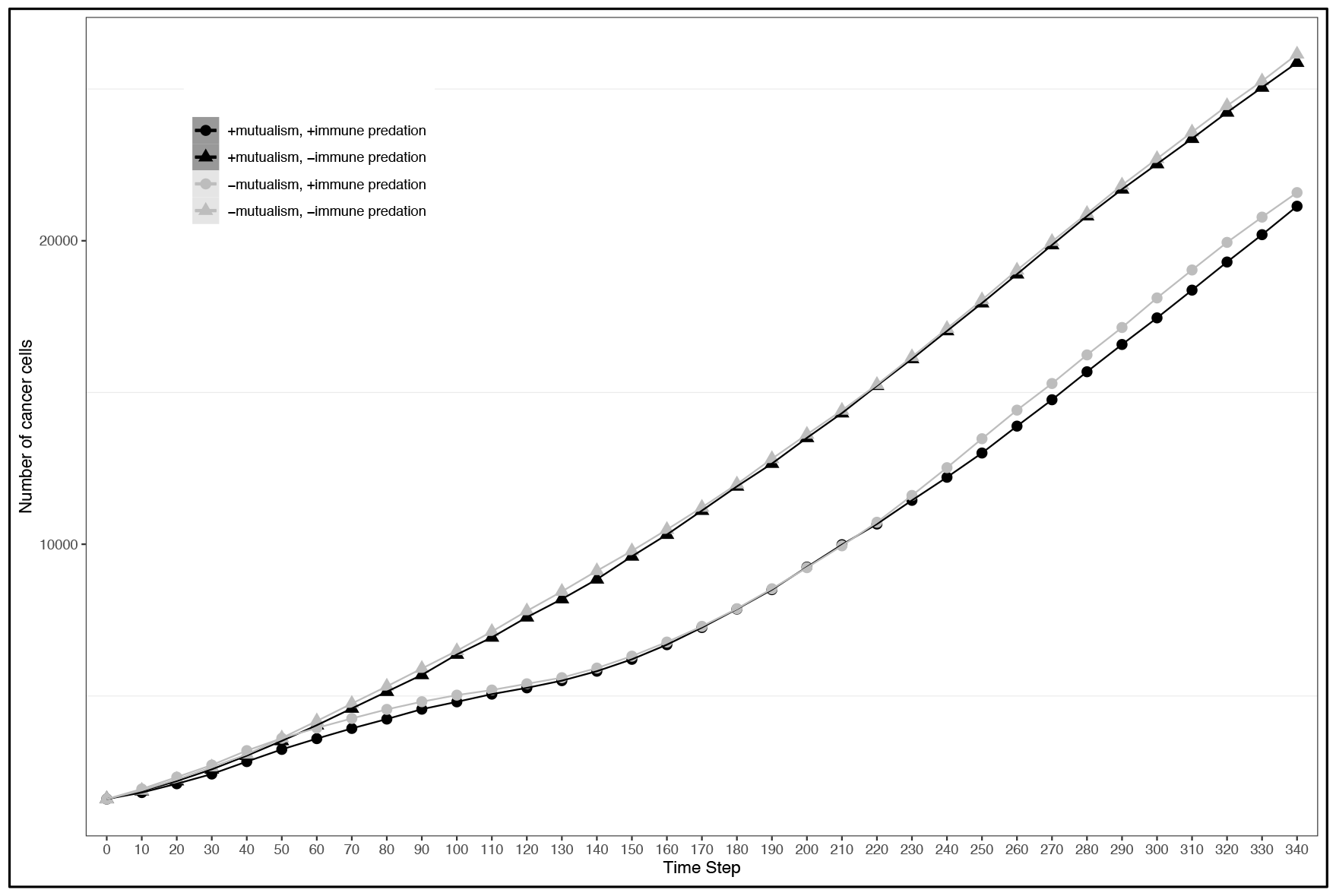
Number of cancer cells depending on different conditions, the division rate in the simulation is 0.09.

**SuppFig2:**
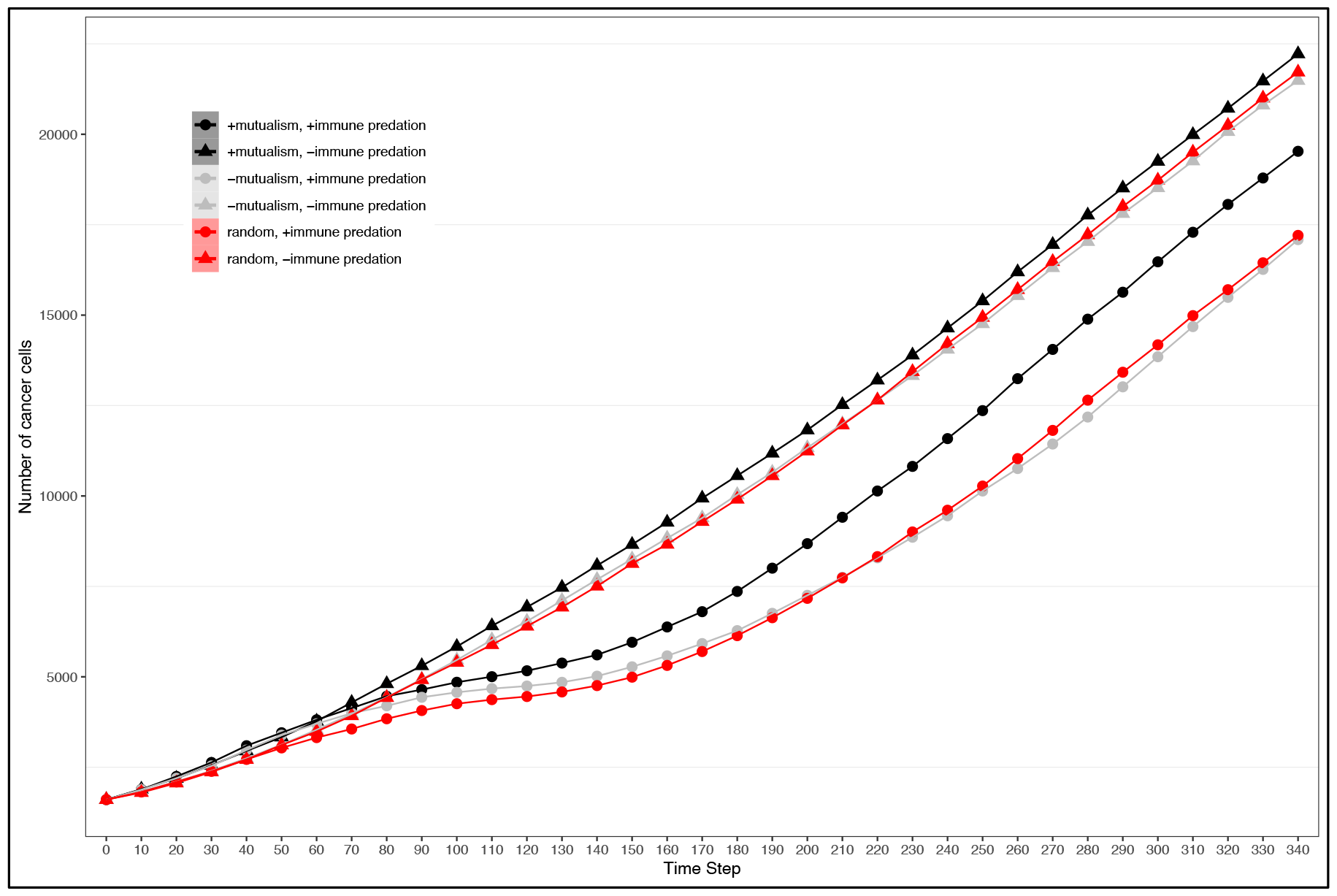
Number of cancer cells for random mutualism, the division rate in the simulation is 0.08.

**SuppFig 3:**
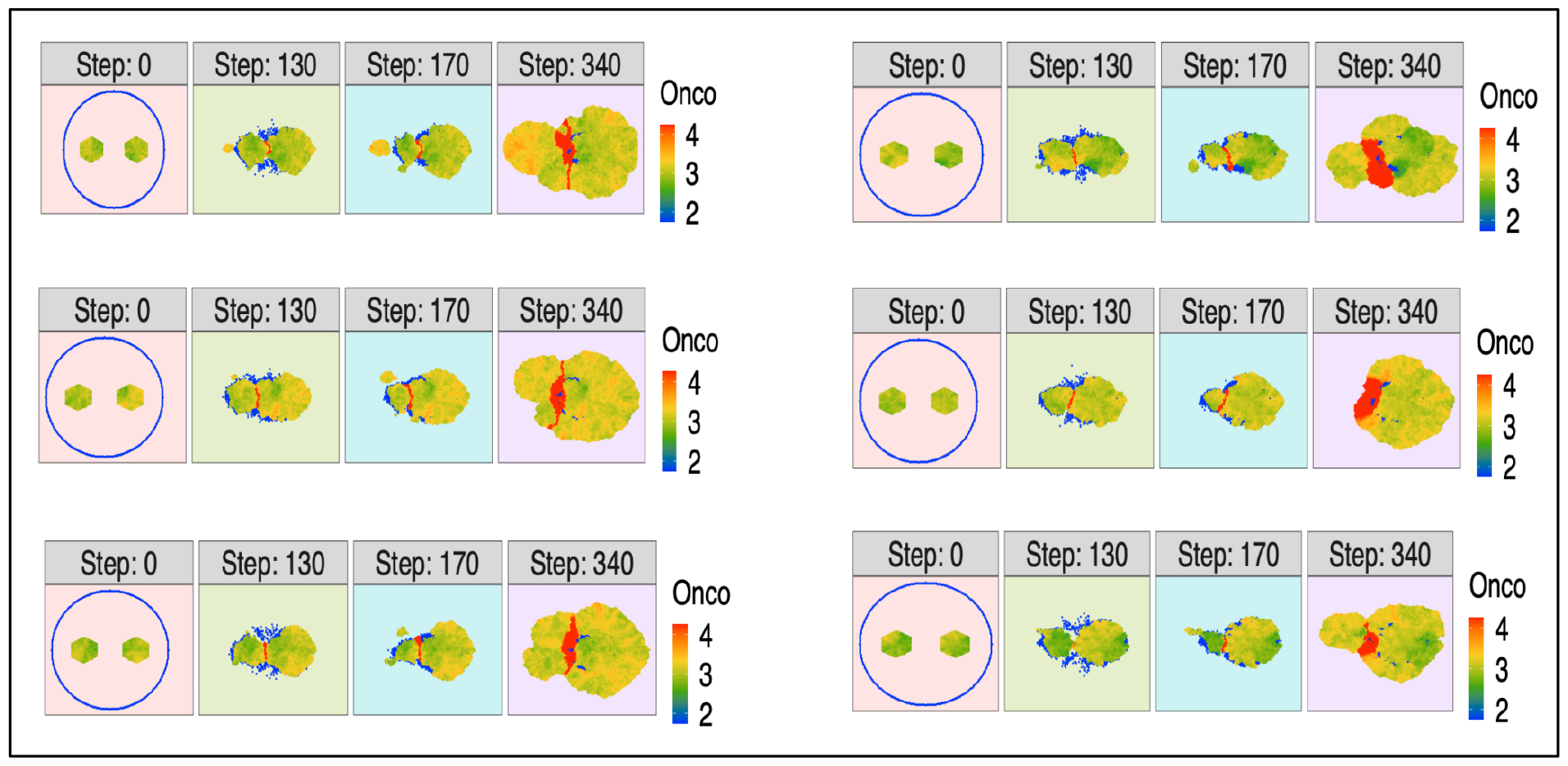
Qualitative comparison of mutualism and no mutualism. On the left simulations supporting our hypothesis, on the right simulations which do not support our hypothesis.

## Author Contributions

Lucie Gourmet conducted the analysis, Parag Mallick provided feedback and reviewed the article. Simon Walker-Samuel also provided feedback and reviewed the article.

## Acknowledgments

I would like to thank the international Alliance for Cancer Early Detection for supporting my PhD and biorender (used to create figures). Moreover, I would like to acknowledge my family, my friends, UCL and Stanford colleagues for their continuous support.

